# A Genetically Encoded Fluorescent Biosensor for Intracellular Measurement of Malonyl-CoA

**DOI:** 10.1101/2024.09.27.615526

**Authors:** Brodie L. Ranzau, Tiffany D. Robinson, Jack M. Scully, Edmund D. Kapelczack, Teagan S. Dean, Tara TeSlaa, Danielle L. Schmitt

## Abstract

Malonyl-CoA is the essential building block of fatty acids and regulates cell function through protein malonylation and allosteric regulation of signaling networks. Accordingly, the production and use of malonyl-CoA is finely tuned by the cellular energy status. Most studies of malonyl-CoA dynamics rely on bulk approaches that take only a snapshot of the average metabolic state of a population of cells, missing out on dynamic changes in malonyl-CoA and fatty acid biosynthesis that could be occurring within a single cell. To overcome this limitation, we have developed a genetically encoded fluorescent protein-based biosensor for malonyl-CoA that can be used to capture malonyl-CoA dynamics in single cells. This biosensor, termed Malibu (malonyl-CoA **i**ntracellular biosensor to understand dynamics), exhibits an excitation-ratiometric change in response to malonyl-CoA binding. We first used Malibu to monitor malonyl-CoA dynamics during inhibition of fatty acid biosynthesis using cerulenin in *E. coli*, observing an increase in Malibu response in a time- and dose-dependent manner. In HeLa cells, we used Malibu to monitor the impact of fatty acid biosynthesis inhibition on malonyl-CoA dynamics in single cells, finding that two inhibitors of fatty acid biosynthesis, cerulenin and orlistat, which inhibit different steps of fatty acid biosynthesis, increase malonyl-CoA levels. Altogether, we have developed a new genetically encoded biosensor for malonyl-CoA, which can be used to sensitively study malonyl-CoA dynamics in single cells, providing an unparalleled view into fatty acid biosynthesis.

## Introduction

Malonyl-CoA is an integral metabolite in central carbon metabolism. Well-known as the primary building block for fatty acids, malonyl-CoA has also been reported to be involved in the downregulation of β-oxidation of acyl-CoAs by the allosteric inhibition of carnitine palmitoyltransferase 1.^1^ Malonyl-CoA is also used for lysine malonylation, which has been linked to inhibited mitochondrial function and β-oxidation along with other manipulations of metabolism.^1–3^ Recently, malonyl-CoA was found to competitively inhibit mammalian target of rapamycin complex 1 (mTORC1).^4^ Due to the importance of malonyl-CoA for metabolism, cell signaling, and post-translational modifications, the direct study of malonyl-CoA dynamics is needed. However, many previous studies were limited in that they relied on bulk approaches that take only a snapshot of the average metabolic state of a population of cells, possibly missing the dynamics of malonyl-CoA in single cells.

To capture the dynamic flux of metabolites in single cell, genetically encoded biosensors have been developed by linking dynamic changes in metabolites with an optical output.^5,6^ Genetically encoded fluorescent protein-based biosensors for lactate, for instance, enabled visualization of real-time glycolytic flux in a population of cells, demonstrating cell-to-cell heterogeneity in glycolysis.^7^ Recently, biosensors for long-chain fatty acyl-CoA esters and acetyl-CoA have been developed to dynamically measure CoA metabolism.^8–10^

While these biosensors have been used to illuminate metabolite dynamics in single cells, we currently lack a fluorescent protein-based biosensor for malonyl-CoA that would allow for the real-time study of malonyl-CoA dynamics in single cells.

Towards illuminating malonyl-CoA dynamics, a bioluminescent-based biosensor for malonyl-CoA was developed. This biosensor, termed FapR-NLuc, used a malonyl-CoA binding bacterial transcription factor, FapR, as a sensing domain.^11^ FapR is a malonyl-CoA sensitive homodimeric transcription factor found in bacteria that modulates transcription of genes encoding lipid biosynthetic enzymes.^12,13^ Importantly, FapR undergoes large conformational changes when binding malonyl-CoA, reflective of its role in transcriptional regulation.^13,14^ In FapR-NLuc, the subunits of a split nanoluciferase (NLuc) were fused to the N- and C-termini of a FapR variant lacking the N-terminal DNA binding domain. Upon malonyl-CoA binding to FapR-Nluc, the split NLuc were brought together, resulting in bioluminescence. This reporter was used to measure subcellular malonyl-CoA dynamics. However, single-cell resolution of malonyl-CoA dynamics could not be achieved with FapR-NLuc, resulting in bulk readouts of malonyl-CoA dynamics.

To overcome the limitations of FapR-NLuc, we sought to develop a sensitive fluorescent protein-based biosensor based on FapR and circularly permutated enhanced green fluorescent protein (cpEGFP). In this study, we report the development of a single-fluorophore excitation-ratiometric biosensor for malonyl-CoA that we termed Malibu (<u>mal</u>onyl-CoA intracellular <u>b</u>iosensor to <u>u</u>nderstand dynamics). We demonstrate that Malibu is specific for malonyl-CoA and can be used to detect changes in malonyl-CoA dynamics in both bacteria and individual mammalian cells. Thus, Malibu enables the sensitive detection of malonyl-CoA in both bulk assays and at the single cell level.

## Results

### Development of Malibu

To develop a fluorescent protein-based malonyl-CoA biosensor, we sought to repurpose the bacterial transcription factor FapR. To maintain the structural dynamics and malonyl-CoA binding of FapR, but to remove the DNA-binding domain, we truncated *Bacillus subtillis* FapR by deleting the N-terminal 43 residues (FapR_Δ43_). Other biosensors based on bacterial transcription factors succeeded when circularly permutated enhanced green fluorescent protein (cpEGFP) was inserted into flexible regions of the protein, so we identified five flexible loops within FapR as candidate insertion sites for cpEGFP.^8,13^ cpEGFP from the calcium biosensor GCaMP6f was inserted into 46 positions within these loops (**Fig 1a**).^15^ We installed N-terminal Ser/Ala/Gly and C-terminal Gly/Thr linkers connecting cpEGFP with the sensing domain, which have been used in an acyl-CoA biosensor as a starting point for developing a malonyl-CoA biosensor.^8^

**Figure 1.**
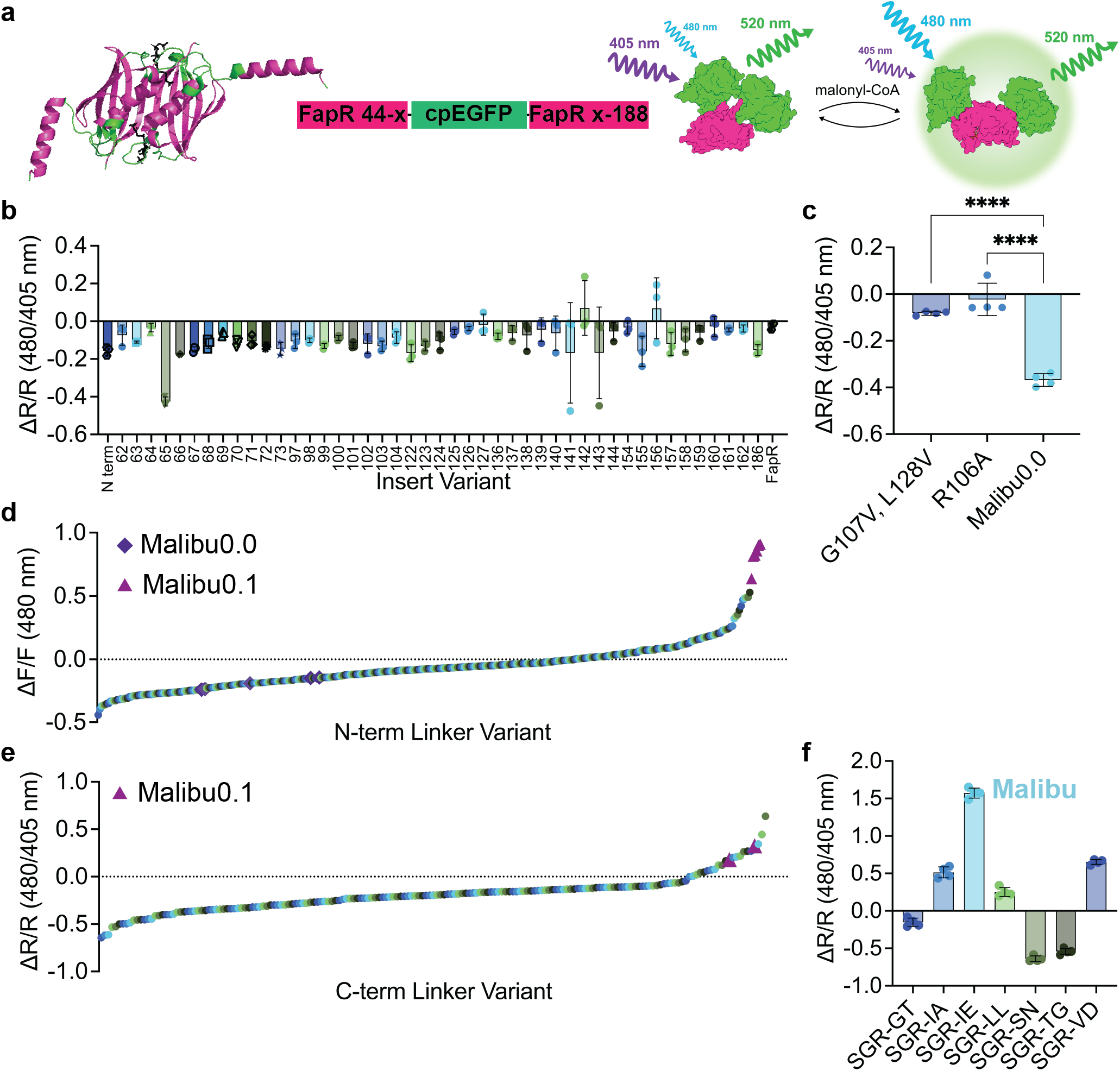
Development of Malibu. **a**, Design and domain layout of Malibu. Crystal structure of FapR, without the N-terminal DNA binding domain, with residues where cpEGFP was inserted highlighted in green and malonyl-CoA in black (PBD ID 2F3X). **b**, Ratio change (ΔR/R) after addition of 500 µM malonyl-CoA of initial Malibu variants in clarified bacterial lysate (n = 3 trials). **c**, Ratio change of Malibu0.0 point mutants expected to minimally bind malonyl-CoA, along with Malibu0.0, treated with 500 µM malonyl-CoA in clarified bacterial lysate (n = 4 trials; p < 0.0001, ordinary one-way ANOVA with Dunnett’s multiple comparisons). **d**, Fluorescence change of N-terminal linker variants screened in clarified bacterial lysate, treated with 500 µM malonyl-CoA. Performance of Malibu0.0 control replicates (purple diamond) and Malibu0.1 (pink triangle) denoted. **e**, Ratio change of C-terminal linker variants screened in clarified bacterial lysate, treated with 450 µM malonyl-CoA. Performance of Malibu0.1 control replicates (pink triangle) denoted. **f**, Ratio change of top hits from C-terminal linker screen in clarified bacterial treated with 450 µM malonyl-CoA (n = 4 trials). For all figures, dot plots show the mean ± SD.

To assess the response of each candidate biosensor, we exposed them to 500 µM malonyl-CoA in bacterial lysate. The change in fluorescence upon malonyl-CoA addition was measured using both 405 nm and 480 nm excitation and 520 nm emission to identify either intensiometric (ΔF/F 480 nm excitation emission) or ratiometric (ΔR/R, 480 nm excitation emission/405 nm excitation emission) reporters, both of which can be developed using cpEGFP.^16,17^ All candidates showed a negative response with 480 nm excitation (ΔF/F 480 nm), as expected with the Ser/Ala/Gly-Gly/Thr linker set that was previously used in a negative-responding acyl-CoA biosensor (**Supplemental Fig 1a**).^8^ When assessing changes in the 480/405 excitation-emission ratio (ΔR/R), only the candidate biosensor with insertion of cpEGFP following Glu 65 showed a consistent change, with ΔR/R of -0.42 ± 0.02 (**Fig 1b)**. We moved forward with this variant, calling it Malibu0.0 (<u>mal</u>onyl-CoA intracellular <u>b</u>iosensor to <u>u</u>nderstand dynamics).

To further confirm that Malibu response is caused by the FapR sensing domain binding to malonyl-CoA, we developed two variants of Malibu that are unable to bind malonyl-CoA. These variants contain either a G107V, L128W double mutation, or a R106A mutation, both of which have been previously reported to not bind malonyl-CoA.^13^ We introduced each of these mutations into Malibu0.0 and observed a response significantly suppressed compared to Malibu0.0 (ΔR/R Malibu0.0 G107V, L128W -0.082 ± 0.009; ΔR/R Malibu0.0 R106A - 0.023 ± 0.069; ΔR/R Malibu0.0 -0.37 ± 0.03, *p* < 0.0001; **Fig 1c**). Thus, the response of Malibu is due to the binding of malonyl-CoA.

Next, we sought to enhance the modest performance of Malibu. The linkers connecting the cpFP with the sensing domain are critical for biosensor function, with mutations in either linker resulting in significant changes to the biosensor response.^18,19^ We began with optimizing the Ser/Ala/Gly linker between the N-terminal FapR fragment and the cpEGFP, focusing on the second and third positions as these residues are close to the chromophore and, in many biosensors, directly influence its fluorescence efficiency and ionization state. A moderate library of variants was generated with these two positions randomly mutated to all amino acids. 379 Malibu variants were randomly selected from this library and tested in bacterial lysate, alongside Malibu0.0 as a control. These variants showed a large range of different responses to malonyl-CoA in lysate, while Malibu0.0 control replicates showed a ΔF/F of -0.19 ± 0.05 and a ΔR/R of -0.39 ± 0.06 (**Fig 1d, Supplemental Fig 1b**). We screened for both intensiometric (ΔF/F) and ratiometric (ΔR/R) changes, finding other variants showed intensiometric responses that ranged from ∼-0.4 to ∼0.8 and ratiometric responses that ranged from ∼-0.6 to ∼0.4. Sequencing the top performing hits from this screen found the top 5 positive responding variants all contained the same Ser/Gly/Arg combination, which in a validation screen had a ΔF/F of 0.81 ± 0.098 (**Supplemental Fig 1c**). Meanwhile, the negative responding variants showed a mixture of combinations, generally enriching for Asn or Ser at either position, with the best performing variant (a Ser/Gly/Asn combination at positions 2 and 3) showing a ΔF/F of -0.28 ± 0.037 in validation screens (**Supplemental Fig 1c)**. Since the Ser/Gly/Arg variant showed the largest absolute response of any Malibu variant thus far, we termed it Malibu0.1 and carried it forward for further optimization.

For the next round of optimization, we mutated both positions of the Gly/Thr linker between cpEGFP and the C-terminal fragment of FapR. We assessed a limited library of 183 random variants alongside Malibu0.1 control, and found that none of the variants performed better than Malibu0.1 when measuring ΔF/F (**Supplemental Fig 1d**). However, when measuring ΔR/R, three variants outperformed Malibu0.1 (**Fig 1e**). Sequencing these variants showed enrichment of a non-polar residue at the first position, with the two best variants further converging on a negatively charged residue at the second position. Validation experiments were performed with the highest performing variants, and confirmed that the Ile/Glu combination works as a ratiometric biosensor and improves the absolute response of Malibu0.1 by 1.5-fold, with a ΔR/R of 1.57 ± 0.07 in bacterial lysate (**Fig 1f**). This variant was selected as our “winner” and is referred to as Malibu.

### In vitro characterization of Malibu

We next characterized Malibu performance *in vitro*. We confirmed that Malibu exhibits a decrease in 405 nm excitation and increase in 480 nm excitation in response to malonyl-CoA, with both excitation peaks contributing to a single emission at 520 nm (**Supplemental Fig 2a**). Malibu exhibited a concentration-dependent change in excitation ratio, reaching a maximum around 1-2 mM malonyl-CoA (**Fig 2a**). Malibu has an affinity for malonyl-CoA of 360 µM (95% confidence interval 240-530 µM). The K_D_ of FapR without the N-terminal DNA binding domain for malonyl-CoA was reported to be 7.1 µM, indicating the insertion of cpEGFP dramatically decreased the affinity of FapR for malonyl-CoA.^11^ A similar trend was observed for the luciferase-based malonyl-CoA biosensor, suggesting that modifications to FapR have impacts on K_D_.^11^

**Figure 2.**
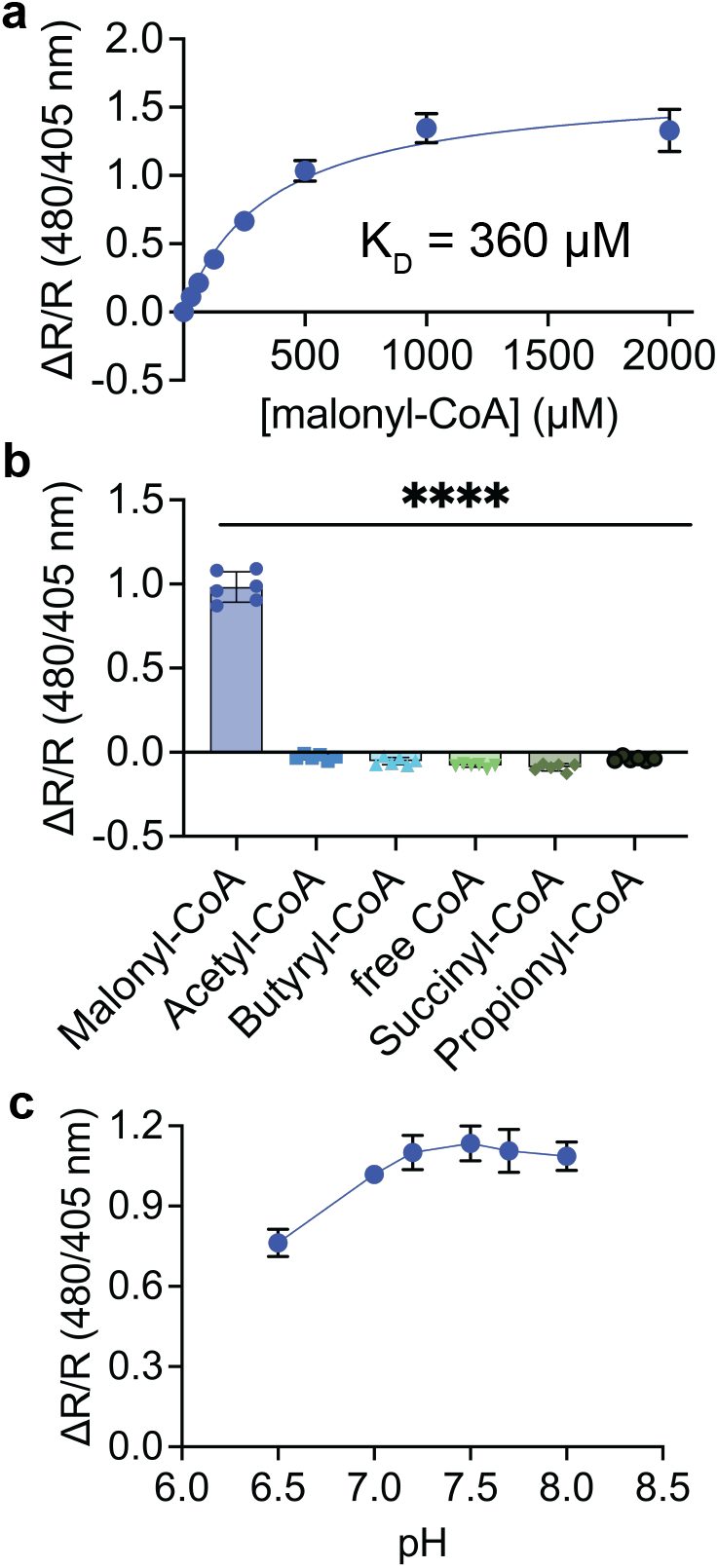
*In vitro* characterization of Malibu. **a**, Dose-response curve for Malibu ratio changes in response to multiple concentrations of malonyl-CoA. Shown are the average ± standard deviation of 18 trials from three independent protein preparations. K_D_ was determined using nonlinear fit. **b**, Selectivity of Malibu towards malonyl-CoA, as measured by ratio change. Malibu was incubated with 500 µM of each respective CoA-containing molecule indicated (6 trials from 2 independent protein preparations; *p* < 0.0001, ordinary one-way ANOVA with Dunnett’s multiple comparison’s test). **c**, pH dependency of Malibu ratio changes in response to 500 µM malonyl-CoA between pH 6.5-8, averaged across 8 trials from two independent protein preparations. For all figures, plots show the mean ± SD.

We next assessed the specificity of Malibu for malonyl-CoA as both purified protein and bacterial lysate. *In vitro* Malibu exhibited a ΔR/R 0.98 ± 0.09 in response to 500 µM malonyl-CoA, but minimally responded to other similar CoAs, including free CoA and acetyl-CoA (*p* ≤ 0.0001; **Fig 2b**). In bacterial lysate, Malibu exhibited similar selectivity towards malonyl-CoA across a range of concentrations (*p* < 0.0001; **Supplemental Fig 2b**).

Finally, we assessed the pH-dependency of Malibu. FP-based biosensors are often pH-dependent.^20,21^ However, ratiometric measurements can correct for this, leading to biosensors with less pH dependency.^22,23^ We found that Malibu in the presence of 500 µM malonyl-CoA exhibited a small increase in ΔR/R from pH 6.5 to pH 7 (ΔR/R 0.76 ± 0.05 to 1.02 ± 0.02) but remained consistent from pH 7-8 (1.02 ± 0.02 to 1.09 ± 0.05; **Fig 2c, Supplemental Fig 2c**). Thus, we have demonstrated that Malibu can sensitively and selectively detect malonyl-CoA, and the ratio change (ΔR/R) does not appear to be pH sensitive at cytoplasmic pH.

### Malibu detects inhibition of fatty acid biosynthesis in bacterial cells

Next, we sought to determine the ability of Malibu to report the intracellular dynamics of malonyl-CoA in bacteria. Inhibition of fatty acid biosynthesis is expected to increase intracellular malonyl-CoA levels, as malonyl-CoA is no longer being actively used for production of new fatty acids. BL21 *E. coli* expressing Malibu were treated with cerulenin, an inhibitor of β-ketoacyl-acyl carrier protein (ACP) synthase I/II (FabB/F), the enzyme responsible for condensing malonyl-ACP with acyl-ACP for nascent fatty acid chain elongation.^24^ Treatment with 4 or 8 µM cerulenin over 2 hours resulted in a significant increase in Malibu ΔR/R compared to treatment with DMSO as measured by plate reader (at 2 hours 4 µM ΔR/R 1.64 ± 0.70; 8 µM ΔR/R 5.03 ± 1.37; DMSO ΔR/R 0.21 ± 0.081; *p* < 0.0001; **Fig 3a**). Under the same conditions, cpEGFP ΔR/R only modestly changed (**Supplemental Fig 3a**). These results are consistent with other findings that inhibiting fatty acid biosynthesis results in an intracellular accumulation of malonyl-CoA in *E. coli*.^25^

**Figure 3.**
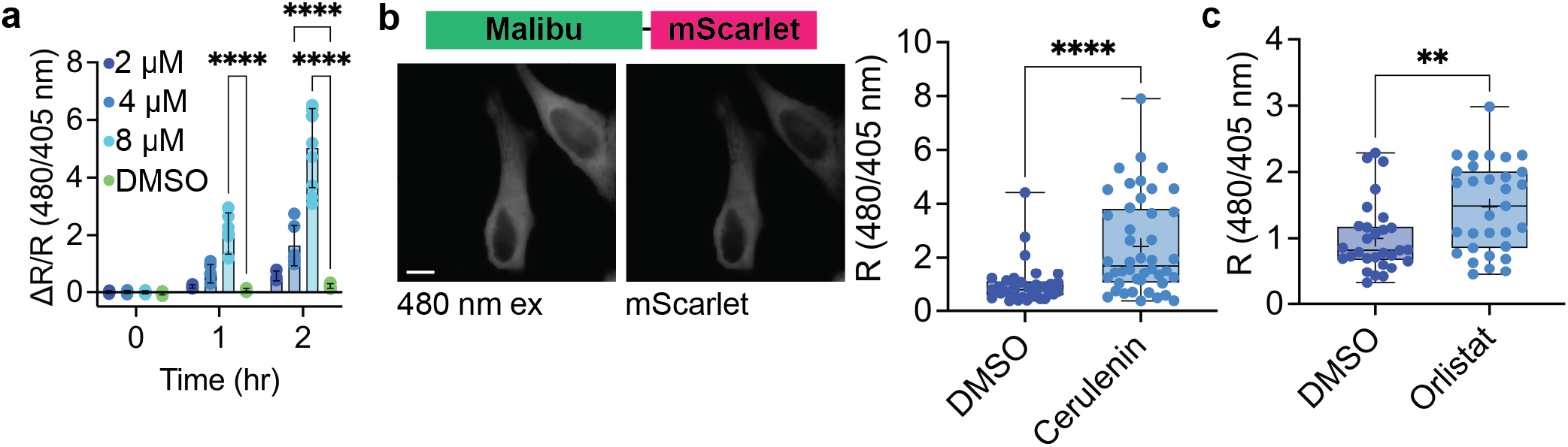
Malibu reports intracellular malonyl-CoA dynamics. **a**, Ratio change of Malibu expressed in BL21 *E. coli* treated with either 2 µM cerulenin (dark blue, n = 8 trials), 4 µM cerulenin (medium blue, n = 8 trials), 8 µM cerulenin (light blue, n = 9 trials), or DMSO (green, n = 9 trials; *p* < 0.0001, 2-way ANOVA with Dunnett’s multiple comparisons test). **b**, Representative images and ratio change of Malibu fused to mScarlet (Malibu-mScarlet) expressed in HeLa cells treated with either DMSO (dark blue, n = 33 cells from 8 experiments) or cerulenin for 4 hr (50 µM, light blue, n = 44 cells from 9 experiments) normalized to mScarlet expression marker (*p* < 0.0001, Mann-Whitney test). **c**, Ratio change of Malibu-mScarlet expressed in HeLa cells treated with either DMSO (dark blue, n = 29 cells from 6 experiments) or orlistat for 12 hr (15 µM, light blue, n = 31 cells from 6 experiments) normalized to mScarlet expression marker (*p* = 0.0074, Mann-Whitney test). For all figures, dot plots show the mean ± SD. For Box-and-Whisker plots, + denotes mean, and min to max is plotted. Scale bar represents 10 µm. Only comparisons with statistical significance are indicated.

### Malibu reports dynamics of fatty acid biosynthesis in mammalian cells

Finally, we used Malibu to capture single-cell dynamics of malonyl-CoA in mammalian cells. We expressed Malibu fused to an mScarlet expression marker (Malibu-mScarlet) in HeLa cells and treated the cells with inhibitors of fatty acid biosynthesis. In mammalian cells, cerulenin inhibits the β-ketoacyl-synthase domain of fatty acid synthase. Following four hours of treatment with cerulenin, we observed a significant increase in Malibu R compared to DMSO control (cerulenin R 2.42 ± 1.82; DMSO R 1 ± 0.79; *p* < 0.0001; **Fig 3b**).

Next, we treated HeLa cells expressing Malibu-mScarlet with orlistat, which inhibits the thioesterase domain of fatty acid synthase.^26^ Following treatment with orlistat for 12 hours, we observed a significant increase in Malibu R compared to DMSO control (orlistat R 1.48 ± 0.68; DMSO R 1 ± 0.53; *p* = 0.0074; **Fig 3c**). To confirm Malibu is faithfully reporting an increase in intracellular malonyl-CoA, we performed LC-MS metabolomics analysis of HeLa cells treated with either cerulenin, orlistat, or DMSO. With both inhibitors of fatty acid biosynthesis, we saw a significant accumulation of malonyl-CoA compared to DMSO control (**Supplemental Fig 3b**). Together, these data demonstrate Malibu can be used to faithfully report the dynamics of malonyl-CoA in single cells.

## Discussion

Genetically encoded biosensors for metabolites are increasingly being developed and used to better understand the dynamic nature of metabolism, as evidenced by the recent reporting of biosensors for glycolytic intermediates and amino acids.^8,9,27–33^ In the present study, we designed a fluorescent protein-based biosensor for malonyl-CoA, Malibu. Consisting of cpEGFP inserted into a truncated bacterial malonyl-CoA binding transcription factor, FapR_Δ43_, Malibu exhibits an excitation-ratiometric change in fluorescence when bound to malonyl-CoA. While a luciferase-based malonyl-CoA biosensor has been previously developed and used to study compartmentalized malonyl-CoA dynamics, the limited sensitivity of the reporter meant only bulk assays could be used to study malonyl-CoA. Using Malibu, we could study inhibition of fatty acid biosynthesis at the single-cell level, with Malibu reporting the accumulation of malonyl-CoA upon inhibition of fatty acid biosynthesis. Furthermore, Malibu exhibited robust selectivity towards malonyl-CoA, providing an advantage over other sensors for CoA derivatives, which suffer from some degree of non-specificity.^8,9^ While Malibu reported an approximate 1.5-fold dynamic range in cells, future engineering efforts will focus on improving the dynamic range of Malibu for the enhanced detection of more subtle malonyl-CoA changes, such as changes in subcellular malonyl-CoA levels.

As we improved Malibu performance, we observed the importance of the linker sequence in determining the response. Initially, we used a Ser/Ala/Gly-Gly/Thr linker combination from a long-chain acyl-CoA biosensor, which resulted in Malibu having a negative response, consistent with response observed in the long-chain acyl-CoA biosensor.^8^ Through random mutagenesis of the N-terminal linker, we identified a Ser/Gly/Arg-Gly/Thr linker combination that flipped the response of Malibu to be a positive, intensiometric response. Further mutagenesis of the C-terminal linkers resulted in no improvement of the intensiometric response, but did reveal a Ser/Gly/Arg-Ile/Glu linker combination that resulted in a positive ratiometric response. With just four mutations, we were able to flip the excitation properties of Malibu.

Analysis of the structure of eLACCO1, a lactate biosensor, enabled the proposed mechanism that amino acids within the linker region interact with the chromophore of cpEGFP to stabilize either the protonated or deprotonated form of the chromophore.^27^ In the case of Malibu, the non-polar Ile residue in the C-terminal linker is likely stabilizing the protonated form of the chromophore, resulting in efficient excitation by 405 nm light. Upon malonyl-CoA binding, conformational changes in the FapR sensing domain may position the Arg residue in the N-terminal linker to better interact with the chromophore, allowing the positive charge to stabilize the deprotonated form, which excites with 480 nm light. This malonyl-CoA-dependent handoff between linker residues likely results in the ratiometric response we observe with Malibu. Future efforts will involve structural studies of Malibu to better understand the biophysical changes within the biosensor and how structural changes in FapR propagate to cpEGFP.

Malibu exhibited an *in vitro* K_D_ of 360 µM, which in our studies was sufficient to measure changes in malonyl-CoA in both bacteria and human cells. The reported concentration of malonyl-CoA in bacteria ranges from 4-90 µM, which suggests that in bacteria, Malibu might not be capturing the full dynamics of malonyl-CoA.^34^ Malonyl-CoA is reported to function as an ATP-competitive inhibitor of mTORC1, with an IC_50_ of 230 µM for human mTORC1 and 334 µM for yeast mTORC1.^4^ This suggests that Malibu is likely to be responsive to concentrations of malonyl-CoA in cells relevant to physiological regulation of fatty acid biosynthesis and metabolic signaling in mammalian cells. Nonetheless, future efforts will involve modifying the apparent affinity of Malibu for malonyl-CoA to best capture the intracellular dynamics of malonyl-CoA, which is especially important as concentrations of malonyl-CoA could vary across subcellular compartments.

In this work, we studied the impact of inhibition of fatty acid biosynthesis in both bacteria and HeLa cells, finding that inhibition increases malonyl-CoA levels, as revealed by increased Malibu ΔR/R. We observe that our measurements of malonyl-CoA in HeLa cells have a large variability in response (**Fig 3b-c**). Similar studies using biosensors to measure metabolic regulation have also had large variability in response and observed significant cell-to-cell variations.^35,36^ As this work was done using an unsynchronized cell population, this variability in response could be due to cell-to-cell differences in metabolic state. Other studies of malonyl-CoA dynamics using luciferase-based malonyl-CoA biosensors highlighted compartmentalized dynamics of malonyl-CoA, finding that orlistat increased malonyl-CoA levels in the cytoplasm and nucleus, but not other locations studied. While not investigated in this work, a potential future application for Malibu is the study of compartmentalized malonyl-CoA dynamics. Furthermore, as the development of red-shifted metabolite biosensors and kinase activity reporters advance, Malibu could be multiplexed with these reporters to provide a more dynamic picture of metabolic regulation in single cells.^7,22,37,38^

In summary, we have generated a genetically encoded fluorescent protein-based malonyl-CoA biosensor, which we have used to study dynamic changes in malonyl-CoA in single cells. Using this biosensor, the study of malonyl-CoA and fatty acid regulation in single cells is now possible, which will result in a greater understanding of metabolic regulation.

## Material and Methods

### Plasmids

All primers used are available in Supplemental Table 1. Primers were obtained from IDT or Eton Bioscience. FapR gene was made by ThermoFisher GeneArt, using FapR sequence from *Bacillus subtillis* (NC_000964.3:1661967-1662533). FapR was truncated (1-43, FapR_Δ43_) to remove the DNA binding domain. Initial designs were cloned by Gibson Assembly (Gibson Assembly HiFi Master Mix, Fisher Scientific Cat# A46628) to insert cpEGFP with SAG-GT linkers into a pRSET vector containing FapR_Δ 43_. The primers for amplifying the backbone, denoted as bb in the primer name, are also denoted by the associated insertion site residue. Primers fwd_insert and rev_insert were used to amplify cpEGFP from GCaMP6f and insert the SAG-GT linkers for all insertion sites except for 67, 99, and 124, which have unique insert primer pairs.

To generate the N-terminal linker library, Golden Gate Assembly was used with primers containing degenerate NNK codons at all positions being mutated. PCR was performed with primers 1-4 with Malibu65 as template, with subsequent DpnI digestion (Thermo Scientific, Cat# FD1703) and PCR cleanup (Thermo Scientific, Cat# K310001), and resulting fragments were subjected to BsmBI digestion and ligation. The ligation was transformed into DH5α cells (Thermo Scientific, Cat# 18265017), which was then diluted into LB for overnight growth. A small aliquot of the transformed cells was plated on LB plates containing 100 µg/mL ampicillin to monitor the number of colony forming units (cfu) produced. >4,000 cfu were detected and the overnight culture was miniprepped (Qiagen, Cat# 27106) to obtain a library of variants.

To generate the C-terminal linker library, Golden Gate Assembly was used with primers containing degenerate NNK codons at the positions being mutated. PCR was performed with primers 8-11 with Malibu0.1 as template. The PCR products were digested with DpnI, cleaned up with a PCR cleanup kit, and subjected to BsmBI digestion and ligation. The library was transformed and isolated following the same protocol as the N-terminal linker library, with >16,000 cfu detected.

Malibu-mScarlet in a pRSET vector was cloned by Gibson Assembly. PCR was performed with primers 9-10 on Malibu in a pRSET vector, and primers 11-12 were used to amplify mScarlet from pEGFP-C1-mScarlet-AMPKα2. Subsequent Gibson Assembly inserted a linker (LQSTGSGNAVGQDTQER), as previously described on the C-terminus of Malibu followed by mScarlet.^33^ Malibu-mScarlet was then inserted into a pcDNA3.1 vector by Gibson Assembly by amplification with primers 15-16, and amplification of the pcDNA3.1 backbone with primers 13-14. QuikChange was performed on this product using primers 17-18 to correct a random mutation that occurred in the product.^39^

To make non-binding mutants of Malibu, QuikChange was used.^39^ The double mutant was made by introducing both mutations individually. The G107V mutation was made with primers 19-20 on template Malibu0.0, and the L128W mutation was made using primers 21-22 using the G107V mutant as template. The R106A non-binding mutant was made using primers 23-24 on Malibu0.0 as template. For all cloning, successful generation was confirmed using either Sanger sequencing (Genewiz) or whole-plasmid long-read sequencing (Plasmidsaurus).

pGP-CMV-GCaMP6f was a gift from Douglas Kim & GENIE Project (Addgene plasmid # 40755;http://n2t.net/addgene:40755; RRID:Addgene_40755). pEGFP-C1-mScarlet-AMPKα2 was a gift from Jin Zhang (Addgene plasmid # 192451; http://n2t.net/addgene:192451; RRID:Addgene_192451).

### Malibu sequence. FapR with cpEGFP in bold

MELSIPELRERIKNVAEKTLEDESGR**NVYIKADKQKNGIKANFKIRHNIEDGGVQLAYHYQQNTPIGDGPVLLP DNHYLSVQSKLSKDPNEKRDHMVLLEFVTAAGITLGMDELYKGGTGGSMVSKGEELFTGVVPILVELDGDVN GHKFSVSGEGEGDATYGKLTLKFICTTGKLPVPWPTLVTTLTYGVQCFSRYPDHMKQHDFFKSAMPEGYIQE RTIFFKDDGNYKTRAEVKFEGDTLVNRIELKGIDFKEDGNILGHKLEYN**IEVKSLSLDEVIGEIIDLELDDQAISILE IKQEHVFSRNQIARGHHLFAQANSLAVAVIDDELALTASADIRFTRQVKQGERVVAKAKVTAVEKEKGRTVVEVNSYVGEEIVFSGRFDMYRSKHS

### Lysate Screening

For lysate assays screening the initial insertion variants, plasmids were transformed into BL21 *E. coli* cells (Fisher Scientific, Cat# C600003) and plated on LB-ampicillin plates. The next day, individual colonies were inoculated in ZYM-5052 auto-induction media with maintenance antibiotics.^40^ Cultures were grown at 37°C with 250 rpm shaking for 6 hours, then 20°C with 250 rpm shaking for 22 hours. Cultures were pelleted and lysed in B-PER (Thermo Scientific Cat# P178248) containing protease inhibitor (Pierce Protease Inhibitor Tablets, Thermo Scientific Cat# A32963), following the manufacturer’s protocol. The resulting lysate was clarified and the supernatant was used for subsequent fluorescence readings by plate reader (Molecular Devices SpectraMax iD5) in a 96-well assay plate (Genesee Scientific, Cat# 33-755). The plate reader was equipped with SoftMax Pro 7.1 data acquisition software (Molecular Devices). Two wells were setup for each sample, one containing malonyl-CoA at a final concentration of 500 µM (Sigma-Aldrich, Cat# M4263), while the other contained no malonyl-CoA. Fluorescence readings were measured with 480 nm and 405 nm excitation, both with 520 nm emission measured. ΔF/F was calculated by taking the difference between the raw fluorescence readings of the paired wells at the indicated fluorescence wavelength and dividing by the reading from the control well or to the same well prior to malonyl-CoA addition. ΔR/R was calculated similarly, but instead of using the raw fluorescence, a ratio was used; which was calculated by dividing the 480 nm excitation-emission by 405 excitation-emission.

For linker screenings, plasmids were transformed into BL21 cells, and individual colonies were inoculated in ZYM-5052 auto induction media in deep well culture plates (Genesee Scientific, Cat# 27-413), covered with a sterile sealing film (Genesee Scientific, Cat# 12-631), and cultured for the time and temperatures previously described. Cultures were pelleted and lysed, and the resulting supernatant was transferred to a 96-well assay plate. Fluorescence measurements before and after addition of 500 µM or 450 µM malonyl-CoA were taken for the N-terminal linker screen and C-terminal linker screen, respectively.

For lysate specificity assays, clarified lysate was pipetted into a 96-well assay plate, and initial fluorescence was measured at 480 and 405 nm excitation with 520 nm emission. Malonyl-CoA, acetyl-CoA (CoALA Biosciences, Cat# AC01), butyryl-CoA (CoALA Biosciences, Cat# BC01), succinyl-CoA (CoALA Biosciences, Cat# SC01), propionyl-CoA (CoALA Biosciences, Cat# PC01), and free CoA (CoALA Biosciences, Cat# CA02) were added in triplicate at final concentrations of 0.50 mM, 0.25 mM, 0.10 mM, and 0.025 mM, and post-addition fluorescence readings were taken.

### Protein Expression and Purification

A 6xHis tag and TEV cleavage site were added to the N-terminus of Malibu in a pRSET vector, using primers 25-28 and subsequent BsmBI digestion and ligation with T4 DNA ligase. BL21 *E. coli* containing plasmid encoding 6xHis-Malibu with a TEV cleavage site between 6xHis tag and Malibu were grown in OM-I media and induced with IPTG (Goldbio, Cat# I2481C50).^41^ Cells were harvested by centrifugation and mechanically lysed in lysis buffer (50 mM Tris, 150 mM NaCl, 20 mM imidazole, 5% glycerol, pH 8.0). Lysate was clarified by centrifugation. Clarified lysate was incubated with Ni-NTA resin (Thermo Scientific, Cat# 25215) and resin was washed with lysis buffer. Then, the resin was incubated with elution buffer (50 mM Tris, 150 mM NaCl, 500 mM imidazole, 5% glycerol, pH 8.0) before collecting the eluate. The eluate was concentrated in 10 kDa MWCO concentrators (Thermo Scientific Cat# 88528). 0.3 mg of TEV protease was added to concentrated eluate to remove 6xHis tag and sample dialyzed using 10 kDa MWCO dialysis membrane (Thermo Scientific Cat# PI88243) in protein storage buffer (50 mM Tris, 150 mM NaCl, 5% glycerol, pH 8.0). Digested and dialyzed eluate was incubated with Ni-NTA resin which was washed using protein storage buffer. The final flow through and all fractions were analyzed via SDS-PAGE (ThermoFisher, Cat# NP0322BOX) and protein purity was assessed. Purified protein was frozen in liquid N_2_ and stored in -80 °C until use.

### In vitro Malibu assays

In vitro characterization of Malibu was performed in a 96-well plate format using black assay plates. All in vitro assays were conducted with 2 nM Malibu in protein assay buffer (50 mM Tris, 150 mM NaCl, 5% glycerol, variable pH). Assay buffer with pH 7.2 was used for specificity and dose-response assays. Assay buffers with pH of 6.5, 7.0, 7.2, 7.5, 7.7, and 8.0 were used in pH sensitivity assays. For K_D_ calculation, nonlinear fit using Equation 1 was done using GraphPad Prism 10.

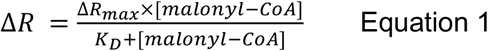

For all assays, fluorescence was read from the top using monochromators set to 405 nm and 480 nm excitation wavelengths and 520 nm emission, detected using an ultra-cooled photomultiplier tube.

### Bacterial fatty acid inhibition assays

BL21 cells were transformed with Malibu and plated overnight on LB-ampicillin plates. Individual colonies were inoculated in ZYM-5052 auto-induction media the subsequent day and incubated at 37°C for 6 hours at 250 rpm, then at 20°C for 22 hours at 250 rpm. Cultures were diluted to OD600 0.60-0.65 in fresh LB media, then pipetted into a clear bottom 96-well assay plate (Thermo Fisher Cat# 152040) along with a media blank to correct for background fluorescence during analysis. Initial fluorescence measurements at 480 nm and 405 nm excitation, both with 520 nm emission, were taken, then Cerulenin was added at final concentrations of 2 µM, 4 µM, and 8 µM along with a DMSO vehicle control. Immediate post-addition fluorescence was measured, and the plate was kept in the plate reader for subsequent readings at 1 and 2 hours post addition. The plate reader temperature was set to 37°C and was programmed to perform 5 seconds of orbital shaking prior to each read.

### Cell culture and transfection

HeLa cells were obtained from ATCC (Fisher Scientific Cat# 50-238-3230). Cells were maintained at 37°C and 5% CO_2_ in Dulbecco’s modified Eagle medium (DMEM, Thermo Scientific Cat# 10569010) containing 4.5 g/L glucose, glutaMAX supplement, sodium pyruvate, 10% fetal bovine serum (Avantor 76419-584) and 100 U/mL penicillin-streptomycin (Pen/Strep, Fisher Scientific Cat# 15-140-122). HeLa cells were routinely checked for mycoplasma using NucBlue (Fisher Scientific Cat #R37605) staining and PCR testing using the PCR Mycoplasma detection kit (Fisher Scientific Cat# AAJ66117AMJ).

Cells were seeded at 150k-200k one day before transfection in 35mm glass bottom dishes (Cellvis, Cat# d35-14-1.5-n). The following day, transfection was carried out in reduced serum medium (Opti-MEM, Thermo Scientific Cat # 31985070) using FuGENE HD (FuGENE, Cat# HD-1000) as previously described.^42^

### Fluorescence imaging and image analysis

Fluorescence microscopy experiments were performed on a Nikon ECLIPSE Ti2 epifluorescence microscope equipped with a CFI60 Plan Fluor 40x Oil Immersion Objective Lens (N.A. 1.3, W.D. 0.2 mm, F.O.V. 25 mm; Nikon), a Spectra III UV, V, B, C, T, Y, R, nIR light engine featuring 380/20, 475/28, and 575/25 LEDs, Custom Spectra III filter sets (440/510/575 and 390/475/555/635/747) mounted in Ti cube polychroic, a Kinetix 22 back-illuminated sCMOS camera (Photometrics), and a stage-top incubator set to 37°C (Tokai Hit). NIS-Elements software (Nikon) was used to control the microscope. Exposure times between 10-100 ms were used.

For HeLa cell fatty acid synthesis inhibition experiments, Orlistat (Alfa Aesar, Cat# J62999-MF) was dissolved in DMSO and added to transfected HeLa cells expressing Malibu-mScarlet at a concentration of 15 µM the day after transfection and incubated for 12 hours prior to imaging. Cerulenin (Millipore Sigma, Cat# C2389) was dissolved in DMSO and added to transfected HeLa cells expressing Malibu-mScarlet at a concentration of 50 µM the day after transfection and incubated for 4 hours prior to imaging. For vehicle control experiments, the same volume of DMSO as inhibitor was added to the cells. Immediately prior to imaging, transfected and treated cells were washed three times with HBSS (ThermoFisher Scientific cat#14065056 supplemented to 1 g/L glucose, 20 mM HEPES, pH 7.3-7.4), the solution was supplemented with the same concentration of inhibitor, and the cells were allowed to incubate at 37°C for 15 minutes before imaging.

Images were taken every 30 seconds for 5 minutes, and the last 2 minutes was averaged for reporting. Image analysis was done as previously described using MATLAB R2024a (MathWorks).^42^ The cpEGFP excitation ratio (Ex488/380) was normalized to mScarlet expression to control cell-to-cell variability in expression, and relative to vehicle control measurements. Cells or dishes that were not fully transfected, dying, or under or over exposed were excluded from analysis.

### LC-MS Metabolomics

HeLa cells, seeded at 150k, were plated 2 days before fatty acid synthesis inhibitor treatment in 6-well plates (Genesee Scientific, Cat# 25-105) and each condition was harvested in triplicate. For Cerulenin treatment, Cerulenin dissolved in DMSO was added to a final concentration of 50uM and incubated for 4 hours before harvesting, and the same volume of DMSO was added to vehicle control wells. For Orlistat treatment, Orlistat dissolved in DMSO was added to a final concentration of 15uM and incubated for 12 hours before harvesting, and the same volume of DMSO was added to vehicle control wells. An additional well was cultured without treatment for packed cell volume measurement during harvesting.

After each treatment incubation period, packed cell volume was measured from the untreated well with PCV tubes (Millipore Sigma, Cat# Z760986). Briefly, the well was washed with HBSS once, incubated with Trypsin (Thermo Scientific, Cat# 12604013), and pipetted into a PCV tube for centrifugation at 2.5k rcf for 1 minute. The treated wells were then washed once with HBSS, and 50 µL of chilled 80% methanol per 1 µL packed cell volume was added, and the plate was incubated on dry ice for 10 minutes. The wells were then scraped with cell scrapers (Genesee Scientific, Cat# 25-270) and the mixture was transferred to microcentrifuge tubes for centrifugation at 16k rcf for 10 minutes at 4°C. The supernatant was then transferred to clean microcentrifuge tubes and stored at -80° until LC-MS analysis.

On the day of LC-MS analysis, samples were again spun at 16k rcf for 10 min at 4°C and supernatant was transferred into LC-MS tubes for analysis. A quadrupole-orbitrap mass spectrometer (Exploris 480 ThermoFisher Scientific) operating in polarity switching mode was coupled to hydrophilic interaction liquid chromatography (HILIC) via electrospray ionization. One full scan was performed from m/z 360-1200 at 90,000 resolution. Additional tSIM scans at 90,000 resolution with isolation window of +/-0.7 m/z were run in positive mode for potential malonyl-CoA ions (C21H38N7O19P3S) with AGC target set to standard and Maximum Injection Time Mode set to Auto. Inclusion list included singly charged ions: +H adduct m/z 854.1229, +NH_4_ adduct m/z 871.1494, +K adduct m/z 892.0788, +Na adduct m/z 876.1048, and no adduct m/z 853.1151. ddMS^2^ was also run for these ions with HCD Collision Energies of 30 and 40, orbitrap resolution of 15,000, ACG target set to standard, and Maximum Injection Time Mode set to Auto. LC separation was achieved with a XBridge BEH Amide column (2.1 mm × 150 mm × 2.5 µm particle size, 130 Å pore size; Water, Milford, MA) using a gradient of solvent A (50 mM ammonium acetate, in 95:5 water:acetonitrile) and solvent B (95:5 acetonitrile:water). Flow rate was 0.4 mL/min. The LC gradient was started at 80% B and was ramped to 60% B over 23.25 min. Between 23.25 and 29.25 min the ratio was held at 60% B followed by re-equilibration at 80% B for 15 min until 22.24 min. Autosampler temperature was 4°C, and injection volume was 10 µL. Malonyl-CoA peak identification was verified by running a standard with malonyl-CoA lithium salt (Sigma-Aldrich, Cat# M4263) which resulted in a detectable peak at m/z 854.1229 and RT 10.18 and MS2 fragments characteristic of malonyl-CoA: m/z 428.0, 347.1, 303.1, 136.1, and 99.1. Data was converted to mzxml format with msconvert and ion counts were exported using MAVEN2.^43^

### Statistics and reproducibility

Figure preparation and statistical analysis were performed using GraphPad Prism 10. For comparison of two parametric data sets, Student’s t test was used. Nonparametric tests were done using Mann-Whitney test. For comparing three or more sets of data, ordinary one-way ANOVA or two-way ANOVA followed by multiple comparisons was done. Statistical significance was defined as *p* < 0.05 with a 95% confidence interval. The number of trials, cells analyzed (n cell), number of independent experiments, and statistical tests used are reported in all figure legends. All dot plots shown depict the mean ± standard deviation.

## Associated Content

Supplementary Figures and Tables are available as Associated Content. This includes additional linker screening information, cpEGFP fluorescence changes, excitation-emission sweeps of Malibu, Malibu specificity in bacterial lysate, unnormalized Malibu pH sensitivity, cpEGFP response to cerulenin in *E. coli*, and results of metabolomics studies.

## Data Availability Statement

All data are available within the main manuscript and the Supporting Information. All plasmids generated in the course of this work will be deposited with the non-profit plasmid repository, AddGene.

## Author Information

### Authors

Brodie L. Ranzau—Department of Chemistry and Biochemistry, University of California Los Angeles, Los Angeles, California, United States of America

Tiffany D. Robinson—Department of Chemistry and Biochemistry, University of California Los Angeles, Los Angeles, California, United States of America

Jack M. Scully—Department of Chemistry and Biochemistry, University of California Los Angeles, Los Angeles, California, United States of America

Edmund D. Kapelczack—Department of Molecular and Medical Pharmacology, University of California Los Angeles, Los Angeles, California, United States of America

Teagan S. Dean—Department of Chemistry and Biochemistry, University of California Los Angeles, Los Angeles, California, United States of America

Tara TeSlaa—Department of Molecular and Medical Pharmacology and Molecular Biology Institute, University of California Los Angeles, Los Angeles, California, United States of America

## Author Contributions

D.L.S., J.M.S., and B.L.R. conceived and designed Malibu. B.L.R. developed and tested Malibu and performed all linker screening assays. J.M.S. purified Malibu and performed all *in vitro* experiments. T.D.R. performed experiments with fatty acid inhibitors. T.D.R., E.D.K., and T.D. performed metabolomics. E.D.K., T.D., and T.T. analyzed metabolomics data. D.L.S. and T.T. oversaw all experiments. B.L.R., T.D.R., J.M.S., T.T., and D.L.S. wrote the manuscript. D.L.S. prepared the figures. All authors contributed to the final version of the manuscript.

## Notes

The authors declare they have no competing interests.

## Acknowledgements

We thank Jin Zhang and Michelle Frei for their helpful critique, Peter DePaola for advice and assistance with protein purification, Robert Campbell, Erin Kim, and all members of the Schmitt Lab for their helpful discussion. This work was supported by the National Institutes of Health (1DP2GM154012 to D.L.S. and T32GM007185 to J.M.S.), the Chan Zuckerberg Initiative (MET-000000000151 to T.T. and D.L.S.), the UCLA Faculty Career Development Award (to D.L.S.), and the UCLA Jonsson Comprehensive Cancer Center Strategic Plan Aligned Project Grant (supported in part by the National Institutes of Health P30CA016042 to D.L.S.).

## Notes

### Competing Interest Statement

The authors have declared no competing interest.

## References

(1) Saggerson, D. Malonyl-CoA, a Key Signaling Molecule in Mammalian Cells. Nutrition 2008, 28 (1), 253– 272. 10.1146/annurev.nutr.28.061807.155434.

(2) Zou, L.; Yang, Y.; Wang, Z.; Fu, X.; He, X.; Song, J.; Li, T.; Ma, H.; Yu, T. Lysine Malonylation and Its Links to Metabolism and Diseases. Aging Dis. 2023, 14 (1), 84–98. 10.14336/ad.2022.0711.

(3) Colak, G.; Pougovkina, O.; Dai, L.; Tan, M.; Brinke, H. te; Huang, H.; Cheng, Z.; Park, J.; Wan, X.; Liu, X.; Yue, W. W.; Wanders, R. J. A.; Locasale, J. W.; Lombard, D. B.; Boer, V. C. J. de; Zhao, Y. Proteomic and Biochemical Studies of Lysine Malonylation Suggest Its Malonic Aciduria-Associated Regulatory Role in Mitochondrial Function and Fatty Acid Oxidation [S]. Mol. Cell. Proteom. 2015, 14 (11), 3056–3071. 10.1074/mcp.m115.048850.

(4) Nicastro, R.; Brohée, L.; Alba, J.; Nüchel, J.; Figlia, G.; Kipschull, S.; Gollwitzer, P.; Romero-Pozuelo, J.; Fernandes, S. A.; Lamprakis, A.; Vanni, S.; Teleman, A. A.; Virgilio, C. D.; Demetriades, C. Malonyl-CoA Is a Conserved Endogenous ATP-Competitive MTORC1 Inhibitor. Nat. Cell Biol. 2023, 25 (9), 1303–1318. 10.1038/s41556-023-01198-6.

(5) Greenwald, E. C.; Mehta, S.; Zhang, J. Genetically Encoded Fluorescent Biosensors Illuminate the Spatiotemporal Regulation of Signaling Networks. Chem. Rev. 2018, 118 (24), 11707–11794. 10.1021/acs.chemrev.8b00333.

(6) Choe, M.; Titov, D. V. Genetically Encoded Tools for Measuring and Manipulating Metabolism. Nat. Chem. Biol. 2022, 18 (5), 451–460. 10.1038/s41589-022-01012-8.

(7) Nasu, Y.; Aggarwal, A.; Le, G. N. T.; Vo, C. T.; Kambe, Y.; Wang, X.; Beinlich, F. R. M.; Lee, A. B.; Ram, T. R.; Wang, F.; Gorzo, K. A.; Kamijo, Y.; Boisvert, M.; Nishinami, S.; Kawamura, G.; Ozawa, T.; Toda, H.; Gordon, G. R.; Ge, S.; Hirase, H.; Nedergaard, M.; Paquet, M.-E.; Drobizhev, M.; Podgorski, K.; Campbell, R. E. Lactate Biosensors for Spectrally and Spatially Multiplexed Fluorescence Imaging. Nat. Commun. 2023, 14 (1), 6598. 10.1038/s41467-023-42230-5.

(8) Wang, W.; Wei, Q.; Zhang, J.; Zhang, M.; Wang, C.; Qu, R.; Wang, Y.; Yang, G.; Wang, J. A Ratiometric Fluorescent Biosensor Reveals Dynamic Regulation of Long-Chain Fatty Acyl-CoA Esters Metabolism. Angew. Chem. Int. Ed. 2021, 60 (25), 13996–14004. 10.1002/anie.202101731.

(9) Smith, J. J.; Valentino, T. R.; Ablicki, A. H.; Banerjee, R.; Colligan, A. R.; Eckert, D. M.; Desjardins, G. A.; Diehl, K. L. A Genetically-Encoded Fluorescent Biosensor for Visualization of Acetyl-CoA in Live Cells. bioRxiv 2024, 2023.12.31.573774. 10.1101/2023.12.31.573774.

(10) Wang, W.; Wang, P.; Zhu, L.; Liu, B.; Wei, Q.; Hou, Y.; Li, X.; Hu, Y.; Li, W.; Wang, Y.; Jiang, C.; Yang, G.; Wang, J. An Optimized Fluorescent Biosensor for Monitoring Long-Chain Fatty Acyl-CoAs Metabolism in Vivo. Biosens. Bioelectron. 2024, 247, 115935. 10.1016/j.bios.2023.115935.

(11) Du, Y.; Hu, H.; Pei, X.; Du, K.; Wei, T. Genetically Encoded FapR-NLuc as a Biosensor to Determine Malonyl-CoA in Situ at Subcellular Scales. Bioconjugate Chem. 2019, 30 (3), 826–832. 10.1021/acs.bioconjchem.8b00920.

(12) Schujman, G. E.; Paoletti, L.; Grossman, A. D.; Mendoza, D. de. FapR, a Bacterial Transcription Factor Involved in Global Regulation of Membrane Lipid Biosynthesis. Dev. Cell 2003, 4 (5), 663–672. 10.1016/s1534-5807(03)00123-0.

(13) Schujman, G. E.; Guerin, M.; Buschiazzo, A.; Schaeffer, F.; Llarrull, L. I.; Reh, G.; Vila, A. J.; Alzari, P. M.; Mendoza, D. de. Structural Basis of Lipid Biosynthesis Regulation in Gram-positive Bacteria. EMBO J. 2006, 25 (17), 4074–4083. 10.1038/sj.emboj.7601284.

(14) Albanesi, D.; Reh, G.; Guerin, M. E.; Schaeffer, F.; Debarbouille, M.; Buschiazzo, A.; Schujman, G. E.; Mendoza, D. de; Alzari, P. M. Structural Basis for Feed-Forward Transcriptional Regulation of Membrane Lipid Homeostasis in Staphylococcus Aureus. PLoS Pathog. 2013, 9 (1), e1003108. 10.1371/journal.ppat.1003108.

(15) Chen, T.-W.; Wardill, T. J.; Sun, Y.; Pulver, S. R.; Renninger, S. L.; Baohan, A.; Schreiter, E. R.; Kerr, R. A.; Orger, M. B.; Jayaraman, V.; Looger, L. L.; Svoboda, K.; Kim, D. S. Ultrasensitive Fluorescent Proteins for Imaging Neuronal Activity. Nature 2013, 499 (7458), 295–300. 10.1038/nature12354.

(16) Lobas, M. A.; Tao, R.; Nagai, J.; Kronschläger, M. T.; Borden, P. M.; Marvin, J. S.; Looger, L. L.; Khakh, B. S. A Genetically Encoded Single-Wavelength Sensor for Imaging Cytosolic and Cell Surface ATP. Nat. Commun. 2019, 10 (1), 711. 10.1038/s41467-019-08441-5.

(17) Yaginuma, H.; Okada, Y. Live Cell Imaging of Metabolic Heterogeneity by Quantitative Fluorescent ATP Indicator Protein, QUEEN-37C. bioRxiv 2021. 10.1101/2021.10.08.463131.

(18) Nasu, Y.; Shen, Y.; Kramer, L.; Campbell, R. E. Structure- and Mechanism-Guided Design of Single Fluorescent Protein-Based Biosensors. Nat. Chem. Biol. 2021, 17 (5), 509–518. 10.1038/s41589-020-00718-x.

(19) Chai, F.; Cheng, D.; Nasu, Y.; Terai, T.; Campbell, R. E. Maximizing the Performance of Protein-Based Fluorescent Biosensors. Biochem. Soc. Trans. 2023, 51 (4), 1585–1595. 10.1042/bst20221413.

(20) Tantama, M.; Martínez-François, J. R.; Mongeon, R.; Yellen, G. Imaging Energy Status in Live Cells with a Fluorescent Biosensor of the Intracellular ATP-to-ADP Ratio. Nat. Commun. 2013, 4 (1), 2550. 10.1038/ncomms3550.

(21) Cho, J.-H.; Swanson, C. J.; Chen, J.; Li, A.; Lippert, L. G.; Boye, S. E.; Rose, K.; Sivaramakrishnan, S.; Chuong, C.-M.; Chow, R. H. The GCaMP-R Family of Genetically Encoded Ratiometric Calcium Indicators. ACS Chem. Biol. 2017, 12 (4), 1066–1074. 10.1021/acschembio.6b00883.

(22) Mehta, S.; Zhang, Y.; Roth, R. H.; Zhang, J.; Mo, A.; Tenner, B.; Huganir, R. L.; Zhang, J. Single-Fluorophore Biosensors for Sensitive and Multiplexed Detection of Signalling Activities. Nat. Cell Biol. 2018, 20 (10), 1215–1225. 10.1038/s41556-018-0200-6.

(23) Chen, M.; Sun, T.; Zhong, Y.; Zhou, X.; Zhang, J. A Highly Sensitive Fluorescent Akt Biosensor Reveals Lysosome-Selective Regulation of Lipid Second Messengers and Kinase Activity. ACS Cent. Sci. 2021, 7 (12), 2009–2020. 10.1021/acscentsci.1c00919.

(24) Kawaguchi, A.; Tomoda, H.; Nozoe, S.; ōMura, S.; Okuda, S. Mechanism of Action of Cerulenin on Fatty Acid Synthetase.1. J. Biochem. 1982, 92 (1), 7–12. 10.1093/oxfordjournals.jbchem.a133933.

(25) Davis, M. S.; Solbiati, J.; Cronan, J. E. Overproduction of Acetyl-CoA Carboxylase Activity Increases the Rate of Fatty Acid Biosynthesis in Escherichia Coli *. J. Biol. Chem. 2000, 275 (37), 28593–28598. 10.1074/jbc.m004756200.

(26) Pemble, C. W.; Johnson, L. C.; Kridel, S. J.; Lowther, W. T. Crystal Structure of the Thioesterase Domain of Human Fatty Acid Synthase Inhibited by Orlistat. Nat. Struct. Mol. Biol. 2007, 14 (8), 704–709. 10.1038/nsmb1265.

(27) Nasu, Y.; Murphy-Royal, C.; Wen, Y.; Haidey, J. N.; Molina, R. S.; Aggarwal, A.; Zhang, S.; Kamijo, Y.; Paquet, M.-E.; Podgorski, K.; Drobizhev, M.; Bains, J. S.; Lemieux, M. J.; Gordon, G. R.; Campbell, R. E. A Genetically Encoded Fluorescent Biosensor for Extracellular L-Lactate. Nat. Commun. 2021, 12 (1), 7058. 10.1038/s41467-021-27332-2.

(28) Keller, J. P.; Marvin, J. S.; Lacin, H.; Lemon, W. C.; Shea, J.; Kim, S.; Lee, R. T.; Koyama, M.; Keller, P. J.; Looger, L. L. In Vivo Glucose Imaging in Multiple Model Organisms with an Engineered Single-Wavelength Sensor. Cell Rep. 2021, 35 (12), 109284. 10.1016/j.celrep.2021.109284.

(29) Koberstein, J. N.; Stewart, M. L.; Smith, C. B.; Tarasov, A. I.; Ashcroft, F. M.; Stork, P. J. S.; Goodman, R. H. Monitoring Glycolytic Dynamics in Single Cells Using a Fluorescent Biosensor for Fructose 1,6-Bisphosphate. Proc. Natl. Acad. Sci. 2022, 119 (31), e2204407119. 10.1073/pnas.2204407119.

(30) Davidsen, K.; Marvin, J. S.; Aggarwal, A.; Brown, T. A.; Sullivan, L. B. An Engineered Biosensor Enables Dynamic Aspartate Measurements in Living Cells. eLife 2024, 12. 10.7554/elife.90024.

(31) Hellweg, L.; Pfeifer, M.; Tarnawski, M.; Thing-Teoh, S.; Chang, L.; Bergner, A.; Kress, J.; Hiblot, J.; Wiedmer, T.; Superti-Furga, G.; Reinhardt, J.; Johnsson, K.; Leippe, P. AspSnFR: A Genetically Encoded Biosensor for Real-Time Monitoring of Aspartate in Live Cells. Cell Chem. Biol. 2024. 10.1016/j.chembiol.2024.05.002.

(32) Hario, S.; Le, G. N. T.; Sugimoto, H.; Takahashi-Yamashiro, K.; Nishinami, S.; Toda, H.; Li, S.; Marvin, J. S.; Kuroda, S.; Drobizhev, M.; Terai, T.; Nasu, Y.; Campbell, R. E. High-Performance Genetically Encoded Green Fluorescent Biosensors for Intracellular L-Lactate. ACS Cent. Sci. 2024, 10 (2), 402–416. 10.1021/acscentsci.3c01250.

(33) Marvin, J. S.; Kokotos, A. C.; Kumar, M.; Pulido, C.; Tkachuk, A. N.; Yao, J. S.; Brown, T. A.; Ryan, T. A. IATPSnFR2: A High-Dynamic-Range Fluorescent Sensor for Monitoring Intracellular ATP. Proc. Natl. Acad. Sci. 2024, 121 (21), e2314604121. 10.1073/pnas.2314604121.

(34) Takamura, Y.; Nomura, G. Changes in the Intracellular Concentration of Acetyl-CoA and Malonyl-CoA in Relation to the Carbon and Energy Metabolism of Escherichia Coli K12. Microbiology 1988, 134 (8), 2249– 2253. 10.1099/00221287-134-8-2249.

(35) Kosaisawe, N.; Sparta, B.; Pargett, M.; Teragawa, C. K.; Albeck, J. G. Transient Phases of OXPHOS Inhibitor Resistance Reveal Underlying Metabolic Heterogeneity in Single Cells. Cell Metab. 2021, 33 (3), 649-665.e8. 10.1016/j.cmet.2021.01.014.

(36) Schmitt, D. L.; Curtis, S. D.; Lyons, A. C.; Zhang, J.; Chen, M.; He, C. Y.; Mehta, S.; Shaw, R. J.; Zhang, J. Spatial Regulation of AMPK Signaling Revealed by a Sensitive Kinase Activity Reporter. Nat. Commun. 2022, 13 (1), 3856. 10.1038/s41467-022-31190-x.

(37) Akerboom, J.; Calderón, N. C.; Tian, L.; Wabnig, S.; Prigge, M.; Tolö, J.; Gordus, A.; Orger, M. B.; Severi, K. E.; Macklin, J. J.; Patel, R.; Pulver, S. R.; Wardill, T. J.; Fischer, E.; Schüler, C.; Chen, T.-W.; Sarkisyan, K. S.; Marvin, J. S.; Bargmann, C. I.; Kim, D. S.; Kügler, S.; Lagnado, L.; Hegemann, P.; Gottschalk, A.; Schreiter, E. R.; Looger, L. L. Genetically Encoded Calcium Indicators for Multi-Color Neural Activity Imaging and Combination with Optogenetics. Front. Mol. Neurosci. 2013, 6, 2. 10.3389/fnmol.2013.00002.

(38) Watabe, T.; Terai, K.; Sumiyama, K.; Matsuda, M. Booster, a Red-Shifted Genetically Encoded Förster Resonance Energy Transfer (FRET) Biosensor Compatible with Cyan Fluorescent Protein/Yellow Fluorescent Protein-Based FRET Biosensors and Blue Light-Responsive Optogenetic Tools. ACS Sens. 2020, 5 (3), 719– 730. 10.1021/acssensors.9b01941.

(39) Liu, H.; Naismith, J. H. An Efficient One-Step Site-Directed Deletion, Insertion, Single and Multiple-Site Plasmid Mutagenesis Protocol. BMC Biotechnol. 2008, 8 (1), 91. 10.1186/1472-6750-8-91.

(40) Studier, F. W. Stable Expression Clones and Auto-Induction for Protein Production in E. Coli. Methods Mol. Biol. (Clifton, NJ) 2013, 1091, 17–32. 10.1007/978-1-62703-691-7_2.

(41) Khalilvand, A. B.; Aminzadeh, S.; Sanati, M. H.; Mahboudi, F. Media Optimization for SHuffle T7 Escherichia Coli Expressing SUMO-Lispro Proinsulin by Response Surface Methodology. BMC Biotechnol. 2022, 22 (1), 1. 10.1186/s12896-021-00732-4.

(42) Schmitt, D. L. Imaging Subcellular AMPK Activity Using an Excitation-Ratiometric AMPK Activity Reporter. Curr. Protoc. 2023, 3 (5), e771. 10.1002/cpz1.771.

(43) Seitzer, P.; Bennett, B.; Melamud, E. MAVEN2: An Updated Open-Source Mass Spectrometry Exploration Platform. Metabolites 2022, 12 (8), 684. 10.3390/metabo12080684.

