## Supplemental Info for "A Genetically Encoded Fluorescent Biosensor for Intracellular Measurement of Malonyl-CoA"

**Ranzau et al Supplemental Figure 1**

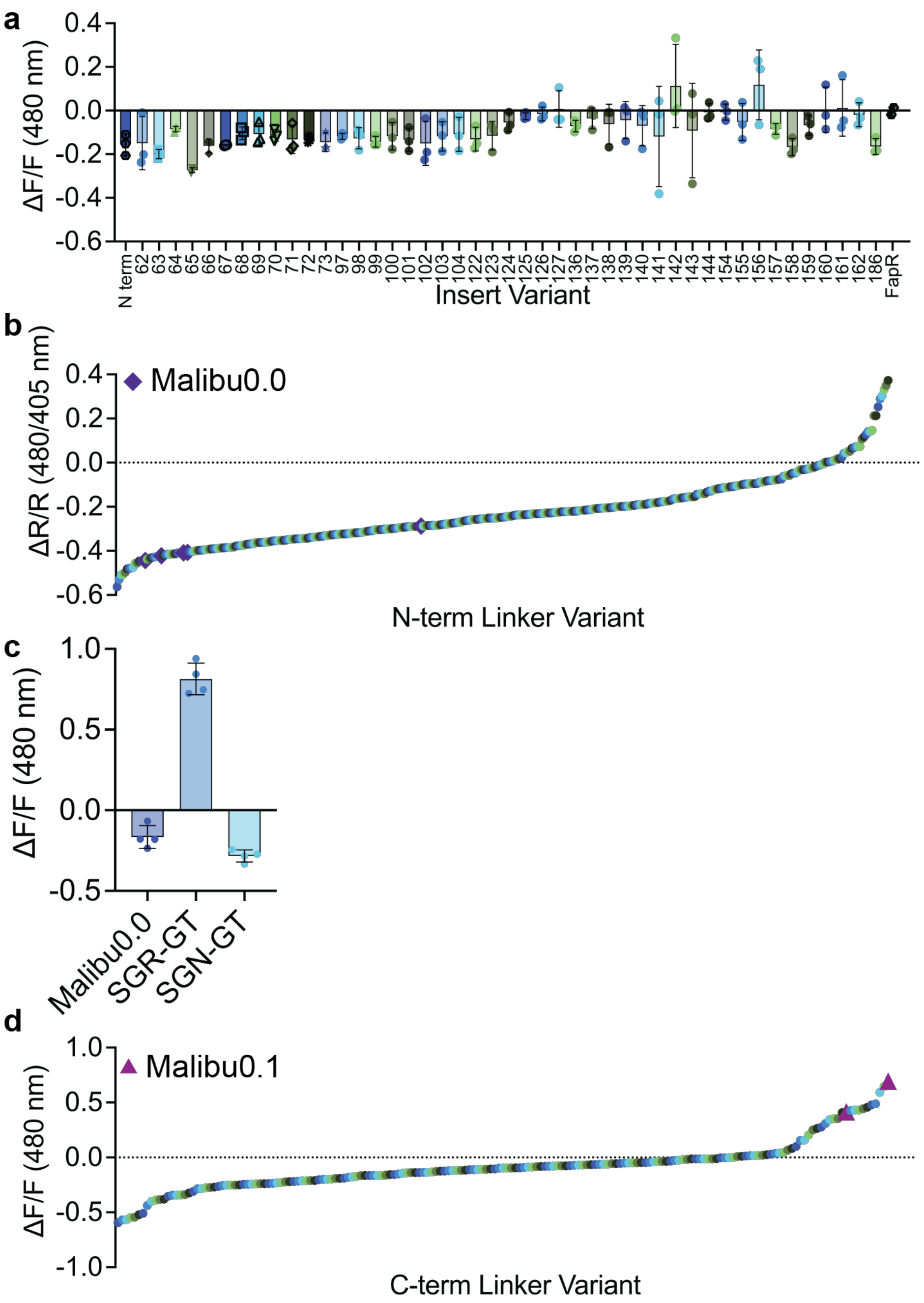


**Supplemental Figure 1. Development of Malibu.**

**a**, Fluorescence change (ΔF/F) after addition of 500 μM malonyl-CoA of initial Malibu variants in clarified bacterial lysate (n = 3 trials).

**b**, Ratio change (ΔR/R) of N-terminal linker variants screened in clarified bacterial lysate, treated with 500 μM malonyl-CoA. Performance of Malibu0.0 (purple diamonds) highlighted.

**c**, Fluorescence change (ΔF/F) of top N-terminal linker variants considered. Malibu0.0 has SAG-GT linkers (n = 4 trials).

**d**, Fluorescence change (ΔF/F) of C-terminal linker variants screened in clarified bacterial lysate, treated with 450 μM malonyl-CoA. Performance of Malibu0.1 (pink triangle) highlighted.

For all figures, dot plots show the mean ± SD.

**Ranzau et al Supplemental Figure 2**

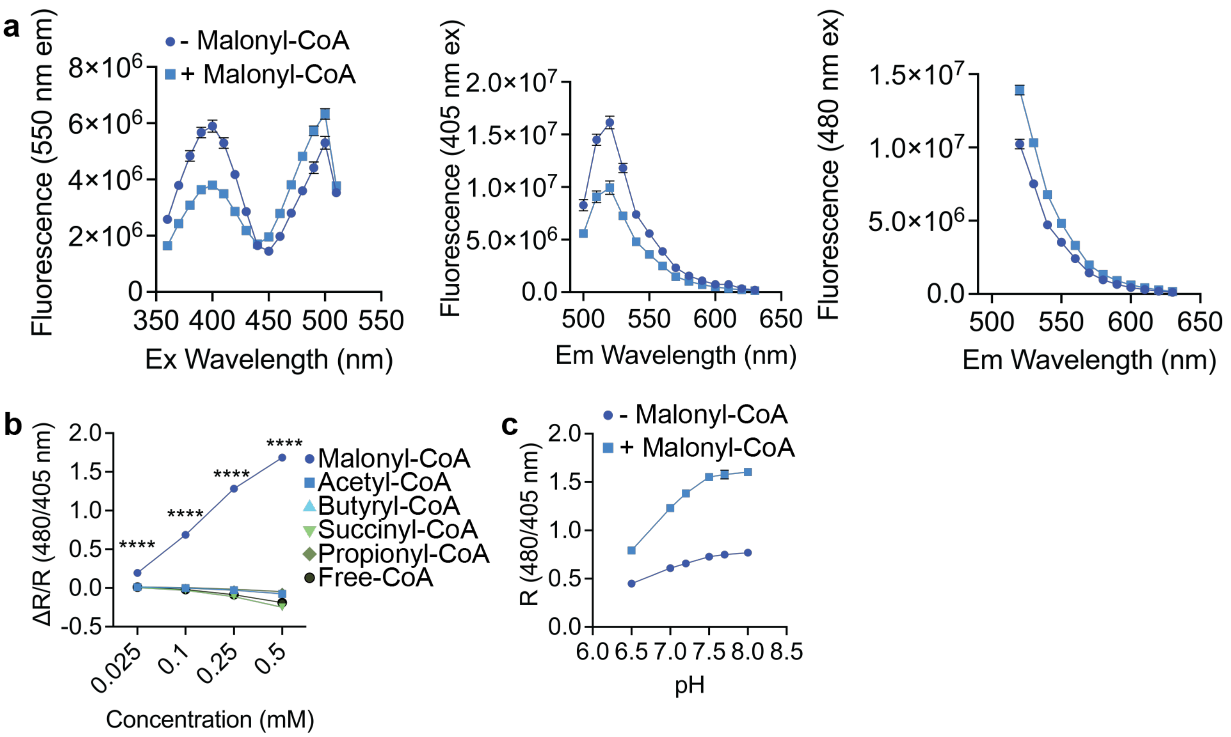


**Supplemental Figure 2. Characterization of Malibu.**

**a,** (left panel) Malibu excitation sweep from 360-510 nm excitation with fluorescence measured at 550 nm emission in the presence of either vehicle (dark blue) or malonyl-CoA (500 μM, light blue). (middle panel) Malibu emission sweep from 500-630 nm with excitation at 405 nm in the presence of either vehicle (dark blue) or malonyl-CoA (500 μM, light blue). (right panel) Malibu emission sweep from 520-630 nm with excitation at 480 nm in the presence of either vehicle (dark blue) or malonyl-CoA (500 μM, light blue). Data represents 4 replicates from 1 protein preparation.

**b**, Selectivity of Malibu towards malonyl-CoA, as measured by ratio change in bacterial lysate. Malibu was incubated with 0.025-0.5 mM of each respective CoA-containing molecule indicated (6 trials; $p < 0.0001$, two way ANOVA with Tukey's multiple comparisons test).

**C**, Unnormalized pH dependency of Malibu ratio changes in response to either vehicle (dark blue) or 500 μM malonyl-CoA (light blue) between pH 6.5-8, averaged across 8 trials from two independent protein preparations.

For all figures, plots show the mean ± SD.

**Ranzau et al Supplemental Figure 3**

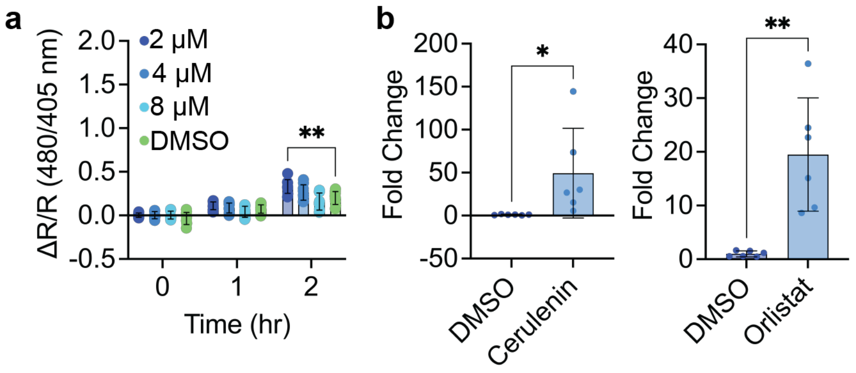


**Supplemental Figure 3. Malibu reports malonyl-CoA dynamics in cells.**

**a**, Ratio change of cpEGFP expressed in BL21 *E. coli* treated with either 2 μM cerulenin (dark blue, n = 8 trials), 4 μM cerulenin (medium blue, n = 8 trials), 8 μM cerulenin (light blue, n = 9 trials), or DMSO (green, n = 9 trials; $p = 0.0052$, 2-way ANOVA with Dunnett's multiple comparisons test).

**b**, (left) Fold change of malonyl-CoA in HeLa cells after treatment with either DMSO (dark blue, n = 6 trials) or cerulenin (50 μM for 4 hr, light blue, n = 6 trials; $p = 0.0465$, unpaired t-test). (right) Fold change of malonyl-CoA in HeLa cells after treatment with either DMSO (dark blue, n = 6 trials) or orlistat (15 μM for 12 hr, light blue, n = 6 trials; $p = 0.0016$, unpaired t-test).

For all figures, dot plots show the mean ± SD.

**Ranzau et al Supplemental Table 1**

| Initial Designs | |
|---|---|
| Primer Number | Sequence 5' to 3' |
| NT_fwd_bb | GGCACAAGCTGGAGTACAACGGAACGGAGCTTTCCATACCTGAACTGAGAGAAAGAATTAAGAACGTGGC |
| NT_rev_bb | GCCTTGATATAGACGTTTCCCGCTGACATGGATCCCCATCGATCCTTATCGTCATCGTCGTACAGATCC |
| 62_fwd_bb | GGCACAAGCTGGAGTACAACGGAACGGAGGACGAAGTGAAGTCCCTGTCACTTGATGAAGTTATCG |
| 62_rev_bb | GCCTTGATATAGACGTTTCCCGCTGAAAGTGTTTTCTCTGCCACGTTCTTAATTCTTTCTCTCAGTTCAG |
| 63_fwd_bb | GGCACAAGCTGGAGTACAACGGAACGGACGAAGTGAAGTCCCTGTCACTTGATGAAGTTATCG |
| 63_rev_bb | GCCTTGATATAGACGTTTCCCGCTGACTCAAGTGTTTTCTCTGCCACGTTCTTAATTCTTTCTCTCAG |
| 64_fwd_bb | GGCACAAGCTGGAGTACAACGGAACGGAAGTGAAGTCCCTGTCACTTGATGAAGTTATCGGAG |
| 64_rev_bb | GCCTTGATATAGACGTTTCCCGCTGAGTCCTCAAGTGTTTTCTCTGCCACGTTCTTAATTCTTTCTCTC |
| 65_fwd_bb | CACAAGCTGGAGTACAACGGAACGGTGAAGTCCCTGTCACTTGATGAAGTTATCGGAGAAATTATTGACC |
| 65_rev_bb | GCCTTGATATAGACGTTTCCCGCTGATTCGTCCTCAAGTGTTTTCTCTGCCACGTTCTTAATTCTTTCTC |
| 66_fwd_bb | GGCACAAGCTGGAGTACAACGGAACGAAGTCCCTGTCACTTGATGAAGTTATCGGAGAAATTATTGACC |
| 66_rev_bb | GCCTTGATATAGACGTTTCCCGCTGACACTTCGTCCTCAAGTGTTTTCTCTGCCACG |
| 67_fwd_bb | TCCCTGTCACTTGATGAAGTTATCGGAGAAATTATTGACCTTG |
| 67_rev_bb | CTTCACTTCGTCCTCAAGTGTTTTCTCTGCCACG |
| 67_fwd_insert | AAAACACTTGAGGACGAAGTGAAGTCAGCGGGAAACGTCTATATCAAGGCCGACAAGCAG |
| 67_rev_insert | CCGATAACTTCATCAAGTGACAGGGACGTTCCGTTGTACTCCAGCTTGTGCCCCAGGATG |
| 68_fwd_bb | GGCACAAGCTGGAGTACAACGGAACGCTGTCACTTGATGAAGTTATCGGAGAAATTATTGACCTTGAGC |
| 68_rev_bb | GCCTTGATATAGACGTTTCCCGCTGAGGACTTCACTTCGTCCTCAAGTGTTTTCTCTGCCACG |
| 69_fwd_bb | GGCACAAGCTGGAGTACAACGGAACGTCACTTGATGAAGTTATCGGAGAAATTATTGACCTTGAGCTGG |
| 69_rev_bb | GCCTTGATATAGACGTTTCCCGCTGACAGGGACTTCACTTCGTCCTCAAGTGTTTTCTCTGCCACG |
| 70_fwd_bb | GGCACAAGCTGGAGTACAACGGAACGCTTGATGAAGTTATCGGAGAAATTATTGACCTTGAGCTGGATG |
| 70_rev_bb | GCCTTGATATAGACGTTTCCCGCTGATGACAGGGACTTCACTTCGTCCTCAAGTGTTTTCTCTGC |
| 71_fwd_bb | GCACAAGCTGGAGTACAACGGAACGGATGAAGTTATCGGAGAAATTATTGACCTTGAGCTGGATGATCAG |
| 71_rev_bb | GCCTTGATATAGACGTTTCCCGCTGAAAGTGACAGGGACTTCACTTCGTCCTCAAGTGTTTTCTCTGC |
| 72_fwd_bb | GGCACAAGCTGGAGTACAACGGAACGGAAGTTATCGGAGAAATTATTGACCTTGAGCTGGATGATCAGGC |
| 72_rev_bb | GCCTTGATATAGACGTTTCCCGCTGAATCAAGTGACAGGGACTTCACTTCGTCCTCAAGTGTTTTCTCTG |

| | |
|---|---|
| 73_fwd_bb | GGCACAAGCTGGAGTACAACGGAACGGTTATCGGAGAAATTATTGACCTTGAGCTGGATGATCAGGC |
| 73_rev_bb | GCCTTGATATAGACGTTTCCCGCTGATTCATCAAGTGACAGGGACTTCACTTCGTCCTCAAGTG |
| 97_fwd_bb | GGCACAAGCTGGAGTACAACGGAACGGTGTTCAGCCGGAATCAGATTGCGAGAGGACACC |
| 97_rev_bb | GCCTTGATATAGACGTTTCCCGCTGAGTGCTCCTGTTTTATTTCTAAAATGGATATCGCCTGATCATCC |
| 98_fwd_bb | GGCACAAGCTGGAGTACAACGGAACGTTCAGCCGGAATCAGATTGCGAGAGGACACC |
| 98_rev_bb | GCCTTGATATAGACGTTTCCCGCTGACACGTGCTCCTGTTTTATTTCTAAAATGGATATCGCCTGATC |
| 99_fwd_bb | AGCCGGAATCAGATTGCGAGAGGACACCAT |
| 99_rev_bb | GAACACGTGCTCCTGTTTTATTTCTAAAATGGATATCGCCTG |
| 99_fwd_insert | GAAATAAAACAGGAGCACGTGTTCTCAGCGGGAAACGTCTATATCAAGGCCGACAAGCAG |
| 99_rev_insert | TGTCCTCTCGCAATCTGATTCCGGCTCGTTCCGTTGTACTCCAGCTTGTGCCCCAGGATG |
| 100_fwd_bb | GGCACAAGCTGGAGTACAACGGAACGCGGAATCAGATTGCGAGAGGACACCATTTATTTGCAC |
| 100_rev_bb | GCCTTGATATAGACGTTTCCCGCTGAGCTGAACACGTGCTCCTGTTTTATTTCTAAAATGGATATCGCC |
| 101_fwd_bb | GGCACAAGCTGGAGTACAACGGAACGAATCAGATTGCGAGAGGACACCATTTATTTGCACAGGC |
| 101_rev_bb | GCCTTGATATAGACGTTTCCCGCTGACCGGCTGAACACGTGCTCCTGTTTTATTTCTAAAATGGATATCG |
| 102_fwd_bb | GGCACAAGCTGGAGTACAACGGAACGCAGATTGCGAGAGGACACCATTTATTTGCACAGGCG |
| 102_rev_bb | GCCTTGATATAGACGTTTCCCGCTGAATTCCGGCTGAACACGTGCTCCTGTTTTATTTCTAAAATGG |
| 103_fwd_bb | GGCACAAGCTGGAGTACAACGGAACGATTGCGAGAGGACACCATTTATTTGCACAGGCGAAC |
| 103_rev_bb | GCCTTGATATAGACGTTTCCCGCTGACTGATTCCGGCTGAACACGTGCTCCTGTTTTATTTCTAAAATGG |
| 104_fwd_bb | GGCACAAGCTGGAGTACAACGGAACGGCGAGAGGACACCATTTATTTGCACAGGCGAACTCTTTGGC |
| 104_rev_bb | GCCTTGATATAGACGTTTCCCGCTGAAATCTGATTCCGGCTGAACACGTGCTCCTG |
| 122_fwd_bb | GGCACAAGCTGGAGTACAACGGAACGGATGACGAGCTGGCGCTGACTGCAAGTGC |
| 122_rev_bb | GCCTTGATATAGACGTTTCCCGCTGAAATGACTGCAACGGCCAAAGAGTTCGCCTGTGC |
| 123_fwd_bb | GGCACAAGCTGGAGTACAACGGAACGGACGAGCTGGCGCTGACTGCAAGTGCAGAC |
| 123_rev_bb | GCCTTGATATAGACGTTTCCCGCTGAATCAATGACTGCAACGGCCAAAGAGTTCGCCTGTGC |
| 124_fwd_bb | GAGCTGGCGCTGACTGCAAGTGCAGACATC |
| 124_rev_bb | GTCATCAATGACTGCAACGGCCAAAGAGTTCG |
| 124_fwd_insert | TTGGCCGTTGCAGTCATTGATGACTCAGCGGGAAACGTCTATATCAAGGCCGACAAGCAG |
| 124_rev_insert | GATGTCTGCACTTGCAGTCAGCGCCAGCTCCGTTCCGTTGTACTCCAGCTTGTGCCCCAG |
| 125_fwd_bb | GGCACAAGCTGGAGTACAACGGAACGCTGGCGCTGACTGCAAGTGCAGACATCCGC |
| 125_rev_bb | GCCTTGATATAGACGTTTCCCGCTGACTCGTCATCAATGACTGCAACGGCCAAAGAGTTCG |

| | |
|---|---|
| 126_fwd_bb | GGCACAAGCTGGAGTACAACGGAACGGCGCTGACTGCAAGTGCAGACATCCGC |
| 126_rev_bb | GCCTTGATATAGACGTTTCCCGCTGACAGCTCGTCATCAATGACTGCAACGGCCAAAGAGTTCG |
| 127_fwd_bb | GGCACAAGCTGGAGTACAACGGAACGCTGACTGCAAGTGCAGACATCCGCTTTACAAGACAGGTAAAGC |
| 127_rev_bb | GCCTTGATATAGACGTTTCCCGCTGACGCCAGCTCGTCATCAATGACTGCAACGGC |
| 136_fwd_bb | GGCACAAGCTGGAGTACAACGGAACGACAAGACAGGTAAAGCAGGGTGAACGTGTCGTAGCAAAAGCG |
| 136_rev_bb | GCCTTGATATAGACGTTTCCCGCTGAAAAGCGGATGTCTGCACTTGCAGTCAGCG |
| 137_fwd_bb | GGCACAAGCTGGAGTACAACGGAACGAGACAGGTAAAGCAGGGTGAACGTGTCGTAGCAAAAGCG |
| 137_rev_bb | GCCTTGATATAGACGTTTCCCGCTGATGTAAAGCGGATGTCTGCACTTGCAGTCAGCGC |
| 138_fwd_bb | GGCACAAGCTGGAGTACAACGGAACGCAGGTAAAGCAGGGTGAACGTGTCGTAGCAAAAGCG |
| 138_rev_bb | GCCTTGATATAGACGTTTCCCGCTGATCTTGTAAAGCGGATGTCTGCACTTGCAGTCAGCGC |
| 139_fwd_bb | GGCACAAGCTGGAGTACAACGGAACGGTAAAGCAGGGTGAACGTGTCGTAGCAAAAGCGAAAGTGACG |
| 139_rev_bb | GCCTTGATATAGACGTTTCCCGCTGACTGTCTTGTAAAGCGGATGTCTGCACTTGCAGTCAGC |
| 140_fwd_bb | GGCACAAGCTGGAGTACAACGGAACGAAGCAGGGTGAACGTGTCGTAGCAAAAGCGAAAGTGACGGC |
| 140_rev_bb | GCCTTGATATAGACGTTTCCCGCTGATACCTGTCTTGTAAAGCGGATGTCTGCACTTGC |
| 141_fwd_bb | GGCACAAGCTGGAGTACAACGGAACGCAGGGTGAACGTGTCGTAGCAAAAGCGAAAGTGACGGC |
| 141_rev_bb | GCCTTGATATAGACGTTTCCCGCTGACTTTACCTGTCTTGTAAAGCGGATGTCTGCACTTGCAGTC |
| 142_fwd_bb | GGCACAAGCTGGAGTACAACGGAACGGGTGAACGTGTCGTAGCAAAAGCGAAAGTGACGGC |
| 142_rev_bb | GCCTTGATATAGACGTTTCCCGCTGACTGCTTTACCTGTCTTGTAAAGCGGATGTCTGCACTTGC |
| 143_fwd_bb | GGCACAAGCTGGAGTACAACGGAACGGAACGTGTCGTAGCAAAAGCGAAAGTGACGGC |
| 143_rev_bb | GCCTTGATATAGACGTTTCCCGCTGAACCCTGCTTTACCTGTCTTGTAAAGCGGATGTCTGC |
| 144_fwd_bb | GGCACAAGCTGGAGTACAACGGAACGCGTGTCGTAGCAAAAGCGAAAGTGACGGC |
| 144_rev_bb | GCCTTGATATAGACGTTTCCCGCTGATTCACCCTGCTTTACCTGTCTTGTAAAGCGGATGTCTGC |
| 154_fwd_bb | GGCACAAGCTGGAGTACAACGGAACGGTCGAAAAAGAAAAAGGAAGAACGGTTGTCGAAGTGAACAGC |
| 154_rev_bb | GCCTTGATATAGACGTTTCCCGCTGAAGCCGTCACTTTCGCTTTTGCTACGACACG |
| 155_fwd_bb | GGCACAAGCTGGAGTACAACGGAACGGAAAAAGAAAAAGGAAGAACGGTTGTCGAAGTGAACAGC |
| 155_rev_bb | GCCTTGATATAGACGTTTCCCGCTGAGACAGCCGTCACTTTCGCTTTTGCTACGACACGTTCACCCTGC |
| 156_fwd_bb | GGCACAAGCTGGAGTACAACGGAACGAAAGAAAAAGGAAGAACGGTTGTCGAAGTGAACAGCTACG |
| 156_rev_bb | GCCTTGATATAGACGTTTCCCGCTGATTCGACAGCCGTCACTTTCGCTTTTGCTACGACACG |
| 157_fwd_bb | GGCACAAGCTGGAGTACAACGGAACGGAAAAAGGAAGAACGGTTGTCGAAGTGAACAGCTACG |

| | |
|---|---|
| 157_rev_bb | GCCTTGATATAGACGTTTCCCGCTGATTTTTCGACAGCCGTCACTTTCGCTTTTGCTACGACACG |
| 158_fwd_bb | GGCACAAGCTGGAGTACAACGGAACGAAAGGAAGAACGGTTGTCGAAGTGAACAGCTACGTTGGC |
| 158_rev_bb | GCCTTGATATAGACGTTTCCCGCTGATTCTTTTTCGACAGCCGTCACTTTCGCTTTTGCTACG |
| 159_fwd_bb | GGCACAAGCTGGAGTACAACGGAACGGGAAGAACGGTTGTCGAAGTGAACAGCTACGTTGGCG |
| 159_rev_bb | GCCTTGATATAGACGTTTCCCGCTGATTTTTCTTTTTCGACAGCCGTCACTTTCGCTTTTGCTACG |
| 160_fwd_bb | GGCACAAGCTGGAGTACAACGGAACGAGAACGGTTGTCGAAGTGAACAGCTACGTTGGCG |
| 160_rev_bb | GCCTTGATATAGACGTTTCCCGCTGATCCTTTTTCTTTTTCGACAGCCGTCACTTTCGCTTTTGC |
| 161_fwd_bb | GGCACAAGCTGGAGTACAACGGAACGACGGTTGTCGAAGTGAACAGCTACGTTGGCG |
| 161_rev_bb | GCCTTGATATAGACGTTTCCCGCTGATCTTCCTTTTTCTTTTTCGACAGCCGTCACTTTCGC |
| 162_fwd_bb | GGCACAAGCTGGAGTACAACGGAACGGTTGTCGAAGTGAACAGCTACGTTGGCGAAG |
| 162_rev_bb | GCCTTGATATAGACGTTTCCCGCTGACGTTCTTCCTTTTTCTTTTTCGACAGCCGTCACTTTCGC |
| 186_fwd_bb | GGCACAAGCTGGAGTACAACGGAACGCATTCATAAGAATTCGAAGCTTGATCCGGCTGCTAACAAAGC |
| 186_rev_bb | CTTGATATAGACGTTTCCCGCTGATTTTGAACGATACATGTCAAAGCGTCCAGAAAAAACAATTTCTTCG |
| fwd_insert | TCAGCGGGAAACGTCTATATCAAGGCCGACAAGC |
| rev_insert | CGTTCCGTTGTACTCCAGCTTGTGCCCCAGG |
| Further developments | |
| Primer Number | Sequence 5' to 3' |
| 1 | AATGCGTCTCTCGAATCANNKNNKAACGTCTATATCAAGGC |
| 2 | TAATCGTCTCATCACCGTTCCGTTGTACTC |
| 3 | AATGCGTCTCTGTGAAGTCCCTGTCACTTGATG |
| 4 | TAATCGTCTCATTCGTCCTCAAGTGTTTTCTC |
| 5 | AATGCGTCTCTGTGAAGTCCCTGTCACTTGATG |
| 6 | TAATCGTCTCATACCAGAACCCCGCATATGTATATCTC |
| 7 | AATGCGTCTCTGGTATGGCTAGCATGACTGG |
| 8 | TAATCGTCTCATCACMNNMNNGTTGTACTCCAGCTTGTGC |
| 9 | GAATTCGAAGCTTGATCCGGCTGCTAACAAAGCCCGAAAGGAAG |
| 10 | GGCCCACCGCGTTGCCGCTGCCGGTGCTCTGCAGTGAATGTTTTGAACGATACATG |
| 11 | GCAACGCGGTGGGCCAGGATACCCAGGAACGCGCCACCATGGTGAGCAAGGGCGAGGCAG |
| 12 | GCCGGATCAAGCTTCGAATTCTTACTTGTACAGCTCGTCCATGCCGCC |
| 13 | GAATTCTGCAGATATCCATCACACTGGCGGCCGCTCG |
| 14 | GATCCTTATCGTCATCGTCGTACAGATCCCGACCCATTTGC |
| 15 | GTACGACGATGACGATAAGGATCCCATGGAGCTTTCCATACCTGAACTGAGAGAAAGAAT |
| 16 | CACCGGCGGCATGGACGAGCTGTACAAGTAAGAATTCTGCAGATATCCATCACACTGGCG |
| 17 | GGCGGCATGGACGAGCTGTACAAGTAAGAATTCTGCAG |

| | |
|---|---|
| 18 | GTCCATGCCGCCGGTGGAGTGGCGGC |
| 19 | CAGATTGCGAGAGTACACCATTTATTTGCACAGGCGAACTC |
| 20 | GTGTACTCTCGCAATCTGATTCCGGCTGAACACGTG |
| 21 | CGTGGACTGCAAGTGCAGACATCCGCTTTACAAGACAGG |
| 22 | CACTTGCAGTCCACGCCAGCTCGTCATCAATGACTGC |
| 23 | GATTGCGGCAGGACACCATTTATTTGCACAGGCGAAC |
| 24 | GTCCTGCCGCAATCTGATTCCGGCTGAACACGTG |
| 25 | TACCGTCTCCCCCATGATGATGATGATGATGAGAACCCATATGTATATCTCCTTCTTAAAG TTAAACAAAATTATTTC |
| 26 | TACCGTCTCCTGGGGGCGGAGAGAATTTGTACTTTCAGGGAGGCGGAGGATCCATGGA GCTTTCCATACCTG |
| 27 | TACCGTCTCCCAGGGTGAACGTGTCGTAGC |
| 28 | TACCGTCTCCCCTGCTTTACCTGTCTTGTAAAGCG |
